# Quantifying the effects of head movement on magnetic resonance spectroscopy estimates of gamma-aminobutyric acid

**DOI:** 10.1101/381699

**Authors:** Edward D. Cui, Richard J. Maddock, Michael H. Buonocore, Meric Ozturk, Jong H. Yoon

**Affiliations:** Depart of Neurosciences, Case Western Reserve University, Cleveland, OH, 44106, United States; Department of Psychiatry and Behavioral Sciences, University of California, Davis, Sacramento, CA, 95817, United States; Department of Radiology, University of California, Davis, Sacramento, CA, 95817, United States; Transportation and Infrastructure Concentration, Price School of Public Policy, University of Southern California, Los Angeles, CA, 90089, United States; Department of Psychiatry and Behavioral Sciences, Stanford University, Stanford, CA, 94305, United States; Palo Alto Veterans Affairs Health Care System, 3801 Miranda Ave., Palo Alto, CA, 94305, United States

**Keywords:** Gamma-aminobutyric acid, Head movement, Magnetic resonance spectroscopy, Neurotransmitters, *in vivo* neuroimaging, Measurement confounds

## Abstract

There has been keen interest in measuring in vivo GABA. However, GABA signal is low and typically measured using techniques vulnerable to the confounding effects of in-scanner head movement. This issue is particularly problematic for clinical studies since it may lead to Type I or II error in testing for group differences. While solutions to mitigate the effects of movement have been proposed, fundamental and largely unexamined issues are the nature and scale of this effect. We developed a method to quantify and characterize head movement during GABA spectroscopy and found that two parameters of movement, displacement and instantaneous movement, were inversely correlated with and accounted for 12.1% and 20.2% variance of GABA estimates respectively. We conclude that head movement can significantly affect GABA measurements and the application of methods to account for movement may improve of GABA measurement accuracy and the detection of true group differences in clinical studies.

## Introduction

As the major inhibitory neurotransmitter in the brain, gamma-aminobutyric acid (GABA) is critical for the regulation of cortical circuit function and is hypothesized to play an important role in a number of neuropsychiatric conditions. GABA signaling enables the synchronization of high frequency oscillatory activity within populations of pyramidal neurons (1, 2). This synchronization is thought to be a fundamental building block for diverse cognitive and perceptual processes supported by cortical circuits (3). Conditions such as schizophrenia (4–6) and depression (7, 8) are thought to involve disruption in GABA production or function. Consequently, there is great interest in measuring in vivo neural GABA levels as this could provide clues to the neural mechanisms underlying information processing (4, 9) and neuropsychiatric conditions (10–12).

Technical developments in proton (^1^H) magnetic resonance spectroscopy (MRS) have led to considerable advances in measuring GABA *in vivo.* A major challenge has been that GABA spectral peaks are obscured by those of other, more abundant metabolites (13). Spectral editing sequences, such as MEscher–GArwood Point RESolved Spectroscopy (MEGA PRESS) have overcome this issue (14, 15) and have been widely adopted by investigators (9, 16–20) to identify altered GABA levels in psychiatric conditions (10, 21, 22). However, the signal for the GABA peak quantified using spectral editing is fairly low (23), raising concerns for its vulnerability to measurement confounds, contributing to false inferences in clinical studies. Thus, there is currently a great need for identifying and accounting for major sources of measurement error.

Head movement during scanning represents one of the most significant confounds for any neuroimaging modality. This is of particular concern in clinical studies in which comparisons are typically made between patient and control groups (24). Significant head movement in control and patient samples could reduce sensitivity to detect true group differences, while group differences in head movement could lead to false conclusions of a group differences (24). In fMRI, methods exist to quantify the amount of head movement during scanning. This quantification can be used to model and statistically correct for the effects of motion on BOLD signal estimations (25) or provide an objective measure by which subjects or portions of scans with excessive movement may be excluded. These methods have allowed investigators to confirm that head movement is a major source of either false positive or false negative findings in fMRI (26–29). The measurement and quantification of head movement is now a basic component of QA in fMRI studies.

Head movement likely has a significant impact on MRS measurements of GABA as well, leading to increased line width, reduced peak intensity, phase incoherence, baseline distortion, and frequency shifts, which can all degrade spectral quality (30). These effects may be particularly problematic when using a frequency selective editing pulse for J-difference spectral editing, as done for GABA measurement with the MEGA-PRESS sequence (31). Methods have been proposed to retrospectively (31–34) and prospectively (30, 35–38) remediate the effects of movement on J-edited GABA measurements. However, fundamental questions as to the scale and the nature of the problem posed by head movement on GABA measurements have been largely unexamined. In addition, the ability to quantify the magnitude of movement occurring during scanning would be helpful at minimum in documenting the magnitude of this confound for a particular study. To address these issues, we developed a method to quantify in-scanner movement during GABA scanning so that we could quantitatively assess the impact head movement has on GABA levels and to make this new tool available for future studies.

## Methods

### Subjects

29 adult subjects free of neurologic or psychiatric illness participated in this study (21 males, 8 females; mean age 22.4 years old, S.D. 3.0 years, range 19 to 32). To confirm the absence of major psychiatric conditions, all subjects underwent the non-patient version of the Structured Clinical Interview for DSM-IV-TR. Additional exclusion criteria were: IQ < 70, drug/alcohol dependence history or abuse in the previous three months, a positive urine drug screen on test day, history of significant head trauma, or any MRI contraindication. Subjects with psychotic first-degree relatives were also excluded. This study was approved by the IRB at University of California, Davis. All subjects provided written informed consent for all procedures.

Subjects were informed about the importance of minimizing head movement and were explicitly instructed to keep their heads as still as possible during scanning. In the scanner, subjects’ heads were secured within the head coil with cushions to minimize head movement.

### MRS data acquisition

We conducted proton MR spectroscopy on the Siemens MAGNETOM Tim Trio 3T MR system using the Siemens 32-channel head RF coil. A 35×30×25 mm voxel was centered on the calcarine sulci bilaterally, with its posterior border approximately 7 mm anterior to the dura. We used a MEGA-PRESS sequence made available by Siemens (Works-in-Progress#529p2) (39) to measure GABA (40). It provides water-suppressed single voxel spectroscopy, with J-difference spectral editing of the GABA C4 resonance at 3.0 ppm. The GABA peak is distinguished from the overlapping creatine singlet (14, 15, 41), through the subtraction of two independently acquired spectra that have GABA C4 J-coupled peaks of opposite polarity. MRS spectral data were acquired using the following parameters: 1500 ms TR; 68 ms TE; 852 ms acquisition duration; 16 step EXOR phase cycling; 1024 data points; 1200 Hz receiver bandwidth; 1.9 ppm and 7.5 ppm spectral edit pulse frequency for on-resonance and off-resonance acquisitions, respectively, taking the conventional approach with the on and off pulses positioned symmetrically about water at 4.7 ppm; 128 acquisitions each for on-resonance and off-resonance spectra; −1.7 ppm delta frequency; spectral edit pulse bandwidth 45 Hz. Including two acquisitions for dynamic equilibrium, total scan time for one run was 6 minutes 27 seconds. We obtained two runs in 6 subjects and 3 runs in 23 subjects.

### Spectroscopy processing and analysis

The on-resonance, off-resonance, and difference spectra were individually zero-filled, phase aligned using the water peak, and apodized using jMRUI v4.0 software. The on and off resonance spectra were frequency aligned on a pair-wise basis by setting the creatine peak to 3.02 ppm. The difference spectra were frequency aligned by setting the midpoint between the peaks of glutamate/glutamine (Glx) to 3.75 ppm. GABA (3.00±0.12ppm in the difference spectra) and creatine (3.02±0.09 ppm in the summed on- and off-resonance spectra) were calculated by peak integration in the frequency domain data using custom software to obtain the GABA/Cr ratios. NAA (2.01±0.14 ppm in the off-resonance spectra) and creatine (3.02±0.09 ppm in the off-resonance spectra) were similarly calculated to obtain the NAA/Cr ratios. GABA and NAA estimates represent the average GABA/Cr and NAA/Cr ratios, respectively, across all runs for each subject.

### Head movement tracking

We monitored head movement by video recording visual markers affixed to the left eye-piece of goggles worn by subjects. These markers were a pair of 3.8 mm × 3.8 mm black squares (inter-centroid distance 4.6mm) printed on a solid white background (Figure 1A). We adapted an in-scanner eye-tracking system of infrared camera and mirrors (Applied Science Laboratory (ASL) EYE-TRAC^®^ 6000), CaptureFlux frame acquisition software acquired video frames of the markers at a sampling rate of 5Hz. Each frame was stored as a 640×480 JPEG image.

**Figure 1.**
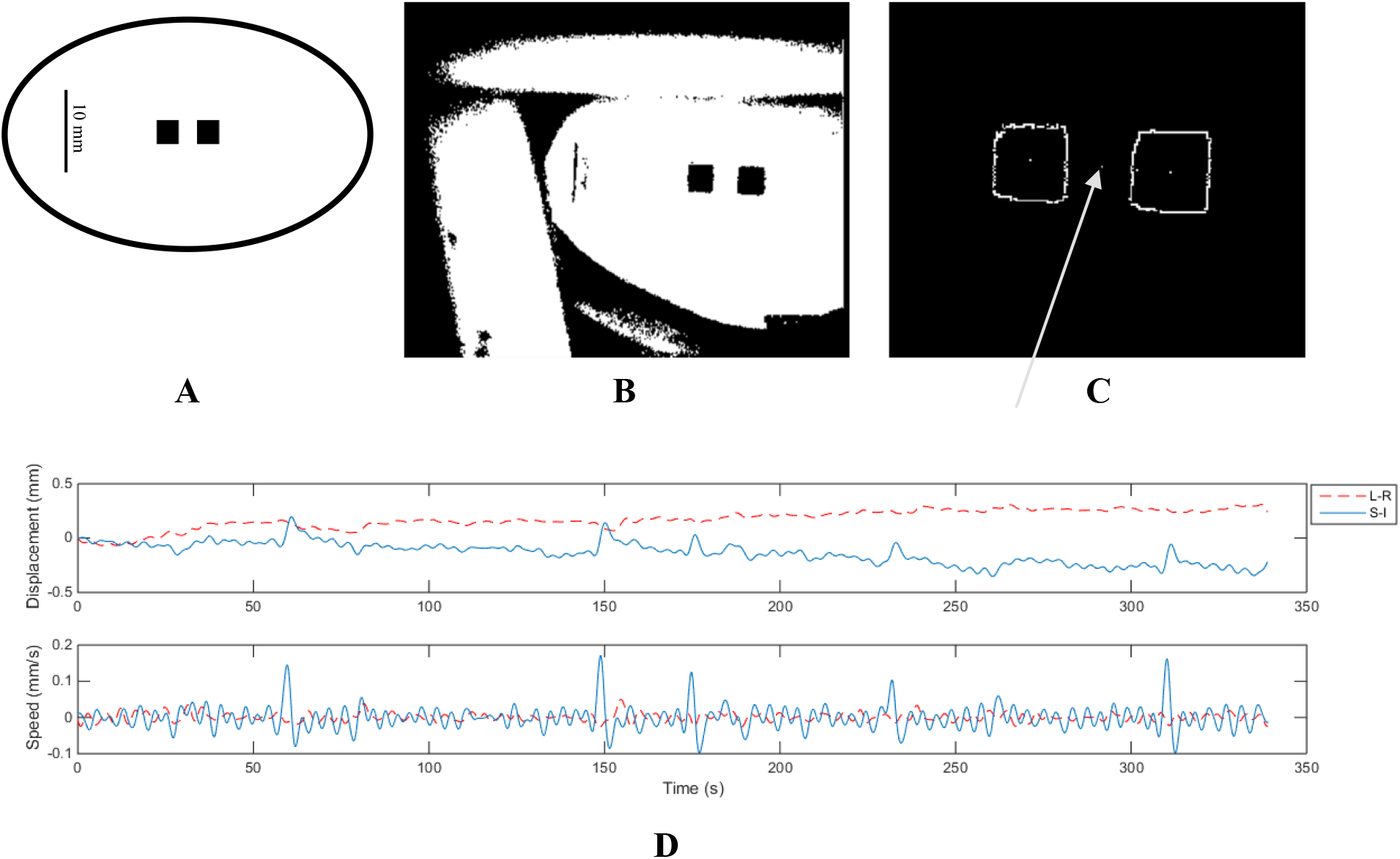
Head movement tracking. **A)** Two-square fiducial marker used in head movement (not drawn to scale); **B)** A binary threshold is selected so that the two squares are clearly distinguishable from each other and from the background; **C)** Centroid of each square is found correctly by the template matching algorithm. The arrow helps the reader to locate the mid-point between the centroids (inter-centroid midpoint) which is the estimated head position at each frame. **D)** Example displacement (top) and velocity (bottom) time series. Legend: “L-R” indicates left-right; “S-I” indicates superior-inferior.

### Head movement data processing

All video frames were processed using scripts developed by the investigators of this study (42). Each frame was converted from grayscale to binary image with thresholds to allow a sufficient separation between the two squares (Figure 1B). Positions of the two squares (Figure 1C) were extracted using a template matching algorithm, Modified Hausdorff Distance (43). The mid-point between the centroids of the squares in the fiducial marker served as the frame-by-frame estimates of head position.

Displacement (mm) was estimated by multiplying the original displacement in pixels with the ratio between the inter-centroid distance in millimeters and in pixels. The resulting time series was filtered with 3^rd^ order Butterworth low pass filter, with 0.25 Hz cutoff frequency.

Since a sMRI was taken prior to the MRS voxel placement, displacement during MRS was referenced to the average position during sMRI. Figure 1D top illustrates an example of the displacement time series. Figure 1D bottom shows velocity, i.e. the first derivative of displacement.

### Head movement quantification

We derived two simple scalar summary metrics to estimate two components of movement: 1) displacement – linear movement away from the original MRS voxel placement at the beginning of the scanning session; 2) speed – moment-by-moment head movement. For each run of MRS, we acquired images of the markers in two spatial dimensions (left-right direction labeled as *x*, and superior-inferior direction labeled as *y*). For each time series of displacement, *D_t_* = (*x_t_, y_t_*), and velocity, 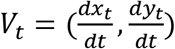:

Magnitude summarizes over spatial dimensions, such that

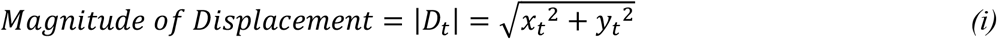

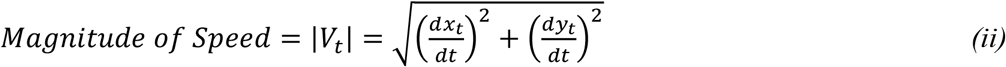

Root Mean Square (RMS) summarizes over time, such that

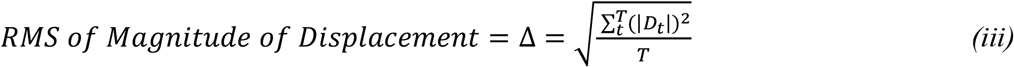

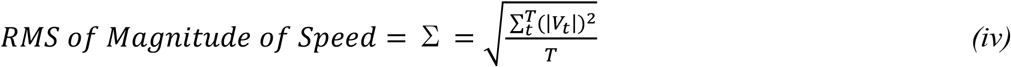

where *t* indexes a single time point, and *T* is the total number of time points. Hence, the two parameters, Δ and Σ, describe average displacement from voxel placement and instantaneous movement, respectively.

### Statistical analysis

All statistical analyses were done in R (44). Model parameters were obtained by maximum likelihood estimation. Plotting was done through ggplot2, an open source graphical package for R (45). We adopted a common practice in fMRI studies of identifying and excluding runs with extreme movement (outliers) (46–49). We assumed that movement had a fixed effect on the normalized GABA estimates and therefore, defined the outlier threshold by pooling all the runs without distinguishing subjects. The outlier threshold was defined by median 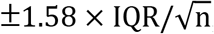, where IQR is the interquartile range, and n is the total number of runs (50). This non-parametric method of defining outliers is advantageous because it does not make assumptions about data distribution (44). Linear regression models were created at subject level, in which averages were taken across runs of each subject.

## Results

### Distributions of movement parameters

Table 1 lists the distribution of each parameter before outlier exclusion, where the “Max” and “Min” are thresholds for outlier exclusion, derived from the non-parametric outlier identification guideline presented in the method section. Following the movement-based data exclusion practice in fMRI, runs with high displacement (Δ > 2.241) or rapid instantaneous movements (Σ > 0.166) are excluded.

**Table 1.** Non-parametric distribution of the dataset.

| Quantile | $\Delta$ (mm) | $\Sigma$ (mm/s) |
| --- | --- | --- |
| *Min* | 0.174 | 0.022 |
| *1<sup>st</sup> Quartile* | 0.672 | 0.047 |
| *Median* | 0.960 | 0.070 |
| *3<sup>rd</sup> Quartile* | 1.363 | 0.098 |
| *Max* | 2.241 | 0.166 |

**Table 2.** Results of linear mixed effect model on repeatability of measurements. *%* change is comparing to the first run.

| Measurements | %Change Run 2<br>(p-value) | %Change Run 3<br>(p-value) |
| --- | --- | --- |
| GABA/Cr | -1.598<br>(0.202) | -5.570<br>(<0.001) |
| Glx/Cr | 1.966<br>(0.347) | 7.067<br>(0.002) |
| NAA/Cr | 0.431<br>(0.419) | 0.302<br>(0.570) |
| Displacement $\Delta$ | 7.791<br>(0.648) | 38.837<br>(0.029) |
| Speed $\Sigma$ | -9.381<br>(0.431) | 5.855<br>(0.622) |

### Spectra with and without movement

Figure 2 shows example spectra from two subjects, one with low movement (Figure 2A, 2B; both Δ and Σ are within the first quartile of the data set) and one with high movement (Figure 2C, 2D, both Δ and Σ are beyond the third quartile of the data set).

**Figure 2.**
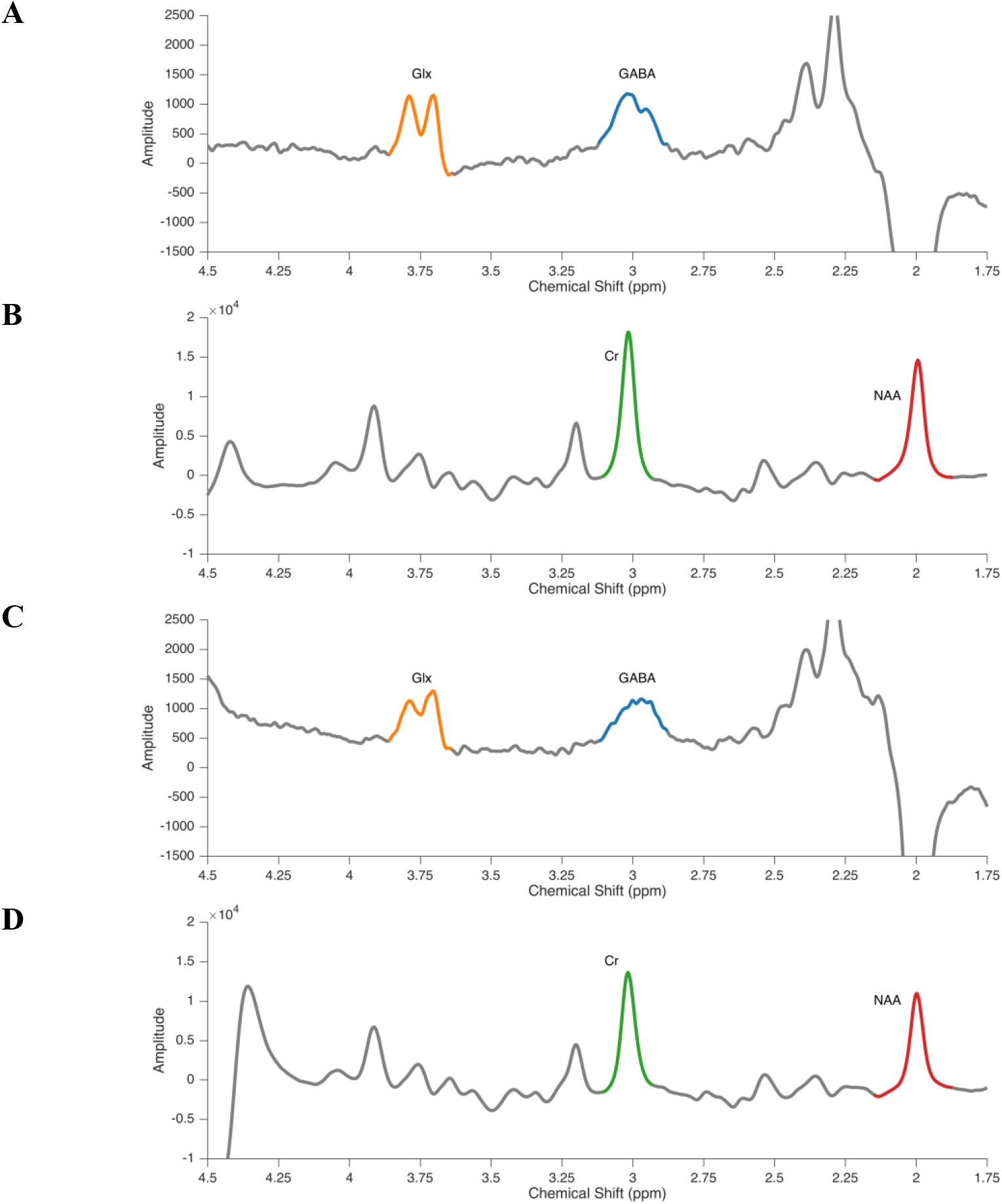
Example spectra data. **A)** Difference and **B)** sum spectra of a run with low movement (displacement Δ = 0.258 mm and scan-to-scan movement Σ = 0.034 mm/s; both within the first quartile of the data set); **C)** Difference and **D)** sum spectra of a run with large movement (Δ = 1.365 mm and Σ = 0.128 mm/s both beyond the third quartile of the data set). Integration ranges are highlighted: GABA in blue, Glx in orange, Cr in green, and NAA in red. Amplitude in arbitrary units. Azimuth unit in parts per million (ppm).

### Associations between movement parameters and neurochemical estimates

To illustrate the effect of individual movement parameters on measurement of GABA, as well as two control compounds, N-acetylaspartate (NAA) and glutamate-glutamine complex (Glx), we constructed the following two models (Figure 3):

**Figure 3.**
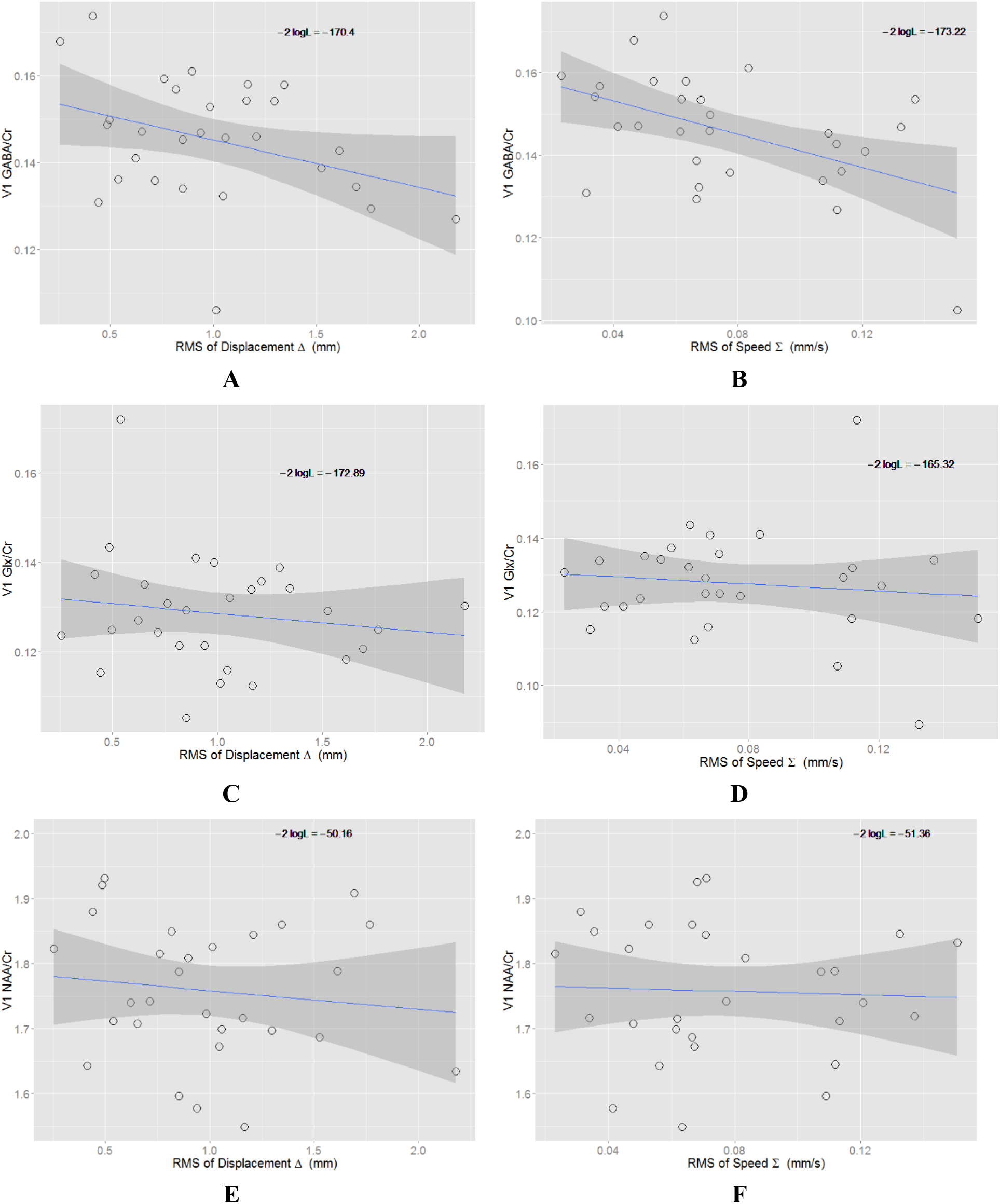
Linear regression models of average head movement parameters and neurochemical measurements of primary visual cortex (V1) in subject level data. N=29 subjects. **A)** GABA/Cr with root-mean-square of displacement (RMS of Displacement Δ) in millimeter (mm); model fit is significant, with P = 0.039; **B)** GABA/Cr with root-mean-square of scan-to-scan movement (RMS of Speed Σ) in millimeter per second (mm/s); model fit is significant, P = 0.002; **C-F)** NAA/Cr and Glx are not significantly impacted by small movements. **C)** Glx/Cr with displacement; **D)** Glx/Cr with speed, P = 0.5386; **E)** NAA/Cr with displacement, P = 0.495; **F)** NAA/Cr with speed, P = 0.814. “-2logL” is an index of fitness of the line to the data, with more negative values indicating better fitness.

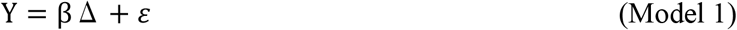

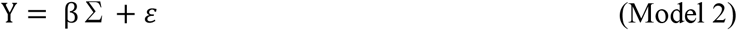

We found that displacement and speed each impacts GABA estimates, but not NAA and Glx (see figure legend for statistical details).

Since displacement from the voxel placement position and instantaneous movement rarely occur separately, we also constructed a model that simultaneously accounts for the effects of displacement and instantaneous movement:

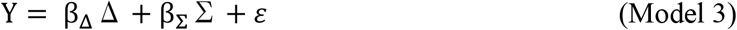

Both displacement and instantaneous movement contribute significantly to GABA estimate (β_Δ_ = –0.011, β_Σ_ = –0.198, z_Δ_ = –1.973, z_Σ_ = –2.885,P_Δ_ = 0.048,P_Σ_ = 0.004,N = 28). ANOVA of the model fit revealed that displacement contributes 12.1% of variance (F = 4.486, P = 0.044) and speed contributed 20.2% of variance (F = 7.476, P = 0.011) in GABA/Cr. Both parameters significantly affect GABA measurement, suggesting that displacement and instantaneous movement are equally important. In contrast, neither displacement nor instantaneous movement contributed significantly to Glx estimate (β_Δ_ = –0.041, β_Σ_ = 0.036, z_Δ_ = –0.775, z_Σ_ = 0.508, P_Δ_ = 0.438, P_Σ_ = 0.612, N = 28) or NAA estimate (β_Δ_ = –0.050, β_Σ_ = 0.006, z_Δ_ = –1.176, z_Σ_ = 0.010,P_Δ_ = 0.240,P_Σ_ = 0.992,N = 28).

## Discussion

We developed a method for quantifying in-scanner head movement and applied it to single voxel J-difference proton spectroscopy of neural GABA. We found that both magnitude of displacement and instantaneous head movement were inversely correlated with GABA estimates. These results have implications for clinical studies relying on between-group comparisons to test for alterations in GABA levels in patient samples.

To the best of our knowledge, this is the first publication describing a method for directly quantifying head movement magnitude during GABA spectroscopy and establishing a quantitative link between head movement and MRS estimates of GABA levels. There have been several previous attempts to account and correct for head movement artifacts on spectral quality of more abundant metabolites, such as water, N-acetylaspartate, and creatine (30, 36, 37). However, as others have noted (31), these methods may not generalize well to GABA, given the possibility that the J-difference spectral editing required for GABA may have unique sensitivity to movement. The absence of a correlation between movement and NAA in our study is consistent with this notion. Of note, the Glx estimates were robust to movement in our study, even though Glx was measured in the same J-edited difference spectra as GABA. However, GABA is obscured by the large creatine peak in unedited spectra, which must be precisely subtracted away in the difference spectra for accurate quantification of GABA. In contrast, the edited Glx peak is not located under an obscuring peak. This suggests that distortion of GABA peak or baseline regions in the difference spectra by imprecise subtraction of the creatine resonance may be a key factor in the effects of motion on GABA estimates. Direct measurement of in-scanner head movement differentiates the present from previous studies and offers a number of advantages that may be useful for future studies. Head movement measures are in standard units of distance, allowing comparisons across populations and studies and the establishment of objective standards for the exclusion of subjects or scan epochs with excessive movement. Individual subjects’ head movement measurements could be used as covariates in statistical models to mitigate the effect of movement on group estimates of GABA levels. From a research standpoint, our method for quantification of movement could allow investigators to assess the impact of motion on the GABA spectra more precisely. Prior studies lacking direct quantification of movement have been limited to qualitative assessments of artifacts in the spectrum presumed to be the results of movement (13, 31).

Our method for quantification of head motion could complement currently available approaches for improving GABA spectra data quality. Magnetic field drift and instability may occur due to a number causes, including head movement, and can induce phase and frequency shifts in the measured spectra. These shifts, in turn, can contribute to increased variability of GABA measurements. Thus, several authors have demonstrated the value of correcting phase and frequency shifts by realigning spectra on a shot-by-shot basis, and rejecting individual problematic subspectra, at post-processing (13, 32–34). Although not used in the current study, this corrective method could be combined with head movement measurements (Figure 1) to identify problematic epochs during scans. High movement epochs for which spectral quality cannot be restored by phase and frequency realignment could be excluded from further analysis.

### Implications for clinical studies

Our results demonstrate that movement can be a source of substantial variance in GABA level estimates. Therefore, not accounting for movement may lead to erroneous conclusions in group comparison studies in two ways. If Group A exhibits significantly greater movement than Group B, this will lower Group A GABA level estimates compared to Group B, and contribute to Type I (false positive) error in group difference in GABA levels. On the other hand, if both Group A and Group B have high movement, the increased variance associated with movement may impede the detection of a true group difference in GABA levels, resulting in a Type II (false negative) error. Overall, the quantification of in scanner movement will improve the detection of true differences in GABA levels by accounting of a significant source of signal variance.

### Limitations

Head movement parameters were estimated in 2-dimensional images, left-right and inferior-superior, leaving out the anterior-posterior dimension. However, in our experience with fMRI experiments in which movement in three dimensions are recorded, movement in the anterior-posterior direction is usually negligible compared to the other directions. On the other hand, an advantage of our method for monitoring head movement in two dimensions is its relative simplicity and ease of use. It requires a single infrared eye-tracking camera, which is readily available in many imaging centers and imposes minimal burden of time and effort to implement. In contrast, most systems monitoring head movement in three dimensions require multiple cameras with complex setups. A method that extracts three dimensional motion parameters from two dimensional image exists (51), but as the authors of this method noted, the sensitivity of the anterior-posterior measurement is compromised if the camera and target are too far from each other, which is most likely the case in the scanner.

It is possible that the visual marker’s movements reflect to some degree non-head-motion related sources, including facial or eyebrow muscle movements, blinking, or scanner vibration. To account for this possibility, we applied a low-pass filter with a cut-off set to exclude high frequency signals from these potentially confounding sources but not to filter out low frequency signals carrying information about head movement. Future endeavors could be aimed at determining the optimal filtering threshold under various scanning conditions.

An important limitation of this study is that data were not acquired in a way that would allow shot by shot phase and frequency correction. Although such corrections are often not included in clinical studies of GABA, this procedure may significantly mitigate movement effects on GABA estimates (31, 32). **Thus, it is unclear to what extent our findings would apply to data sets that have incorporated a shot by shot correction procedure.** In spite of this limitation, our findings demonstrate the utility of a new tool for quantifying intra-scan head movement and determining the extent to which it influences GABA estimates.

### Conclusion

We developed a simple method to measure and quantitatively characterize head movement during GABA MRS scanning. Head movement can be quantified in terms of displacement from the original voxel placement position and instantaneous movement, and both parameters can significantly contribute to variability in GABA estimates. These methods can be readily applied to future studies to limit the effects of in-scanner head movement on estimates of GABA, particularly for clinical studies in which head movement may contribute to erroneous conclusions of group differences in GABA levels.

## Acknowledgments

Research reported in this paper was supported in part by the National Institute of Mental Health of the National Institutes of Health under award number R21MH090475 for JHY. The authors would like to thank Dennis Thompson at UC Davis Imaging Research Center for providing technical support for the head motion tracking method used in this paper.

